# GeneFriends 2021: Updated co-expression databases and tools for human and mouse genes and transcripts

**DOI:** 10.1101/2021.01.10.426125

**Authors:** Priyanka Raina, Inês Lopes, Kasit Chatsirisupachai, Zoya Farooq, João Pedro de Magalhães

## Abstract

Gene co-expression analysis has emerged as a powerful method to provide insights into gene function and regulation. The rapid growth of publicly available RNA-sequencing (RNA-seq) data has created opportunities for researchers to employ this abundant data to help decipher the complexity and biology of genomes. Co-expression networks have proven effective for inferring relationship between the genes, for gene prioritization and for assigning function to poorly annotated genes based on their co-expressed partners. To facilitate such analyses we created previously an online co-expression tool for humans and mice entitled GeneFriends. To continue providing a valuable tool to the scientific community, we have updated the GeneFriends database. Here, we present the latest version of GeneFriends, which includes updated gene and transcript co-expression networks based on RNA-seq data from 46,475 human and 34,322 mouse samples. GeneFriends is freely available at http://www.genefriends.org/

## INTRODUCTION

The advent of RNA sequencing (RNA-seq) technology has revolutionized biological research (Emrich et al. 2007; Lister et al. 2008). With RNA-seq we are able to understand the complexity of transcriptome, which has enabled us to connect the information on our genome with its functional protein expression (Ozsolak and Milos 2011). Moreover, gene co-expression networks provide the potential to identify the gene modules (highly connected sub-networks) that could serve as points for therapeutic interventions (Chen et al. 2008; Cheng et al. 2020). There are many methods available to cluster the genes in a gene co-expression network (see the review; Sipko et al. 2018). One of the widely used network-based approach to predict gene functions is the Guilt by association (GBA) method, GBA works on the principle that genes which tend to co-express with each other are functionally related (Oliver 2000; Molet et al. 2013).

With an increase of more than 2 million RNA-seq samples in SRA/GEO between 2015 and 2020, the number and power of co-expression databases has also consequently increased. (Franz et al. 2018; Wong et al. 2018; Obayashi et al. 2019). To facilitate and promote the usage of co-expression networks, we previously created an online microarray and RNA-seq based co-expression database, entitled GeneFriends (van Dam et al. 2012; vanDam et al. 2015) for human and mouse genes and for human transcripts. GeneFriends has proven successful for gene prioritization and associating function to poorly annotated genes. Studies employing GeneFriends have focused on diverse topics such as estimating tumorigenic index for cancer initiation and progression (Wang et al. 2019), genetic analysis for neurological conditions in humans and mice (Ashbrook et al. 2015), genomics of human metabolic disease (Timmons et al. 2018), development of neuronal subtypes (Memic et al. 2016), genome evolution (Keane et al. 2015), genetics of ageing and complex diseases (Fernandes et al. 2016; Marttila et al. 2020) and cell senescence (Avelar et al. 2020). Therefore, to keep our tool at the forefront of publicly available co-expression databases we have updated the RNA-seq based GeneFriends co-expression database for both human and mouse data. We believe our latest updated version of GeneFriends will be useful for a diverse and large number of researchers to understand the complexity, functions and regulation of the human and mouse genomes. GeneFriends is freely available at http://www.genefriends.org.

## GENEFRIENDS UPDATED CO-EXPRESSION DATABASE

In addition to updating the previous GeneFriends co-expression database for human genes and transcripts (van Dam et al. 2015), we have now added an RNA-seq based co-expression database for mouse genes and transcripts (Figure 1).

**Figure 1.**
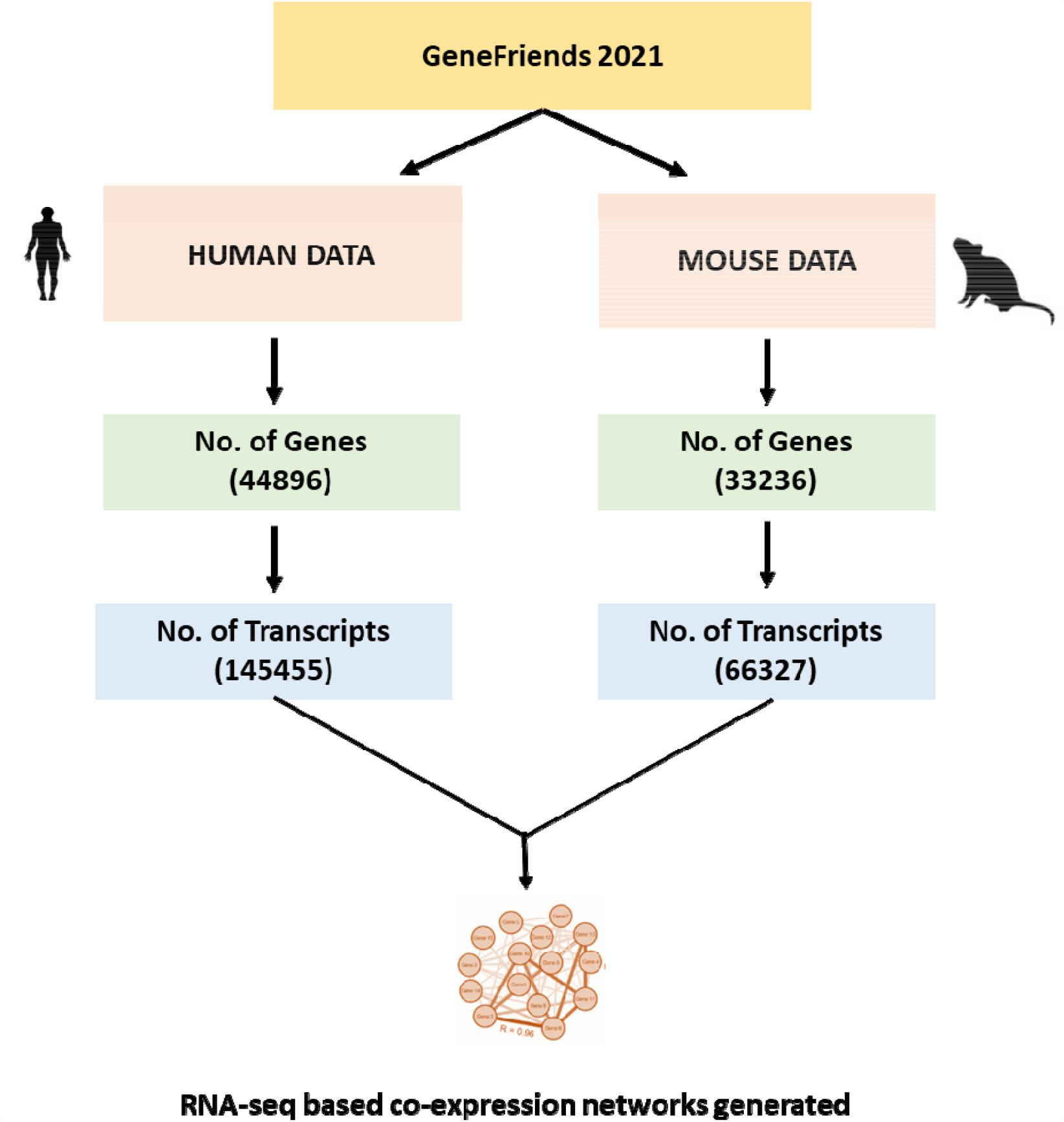
Overview of updated RNA-seq GeneFriends co-expression database for human and mouse genes and transcripts. Human RNA-seq read counts were downloaded from *recount2* database and mouse read counts were obtained from the ARCHS^4^ database.

### Human and Mouse co-expression gene and transcript data

The new human and mouse co-expression databases were constructed from 46,475 and 34,322 RNA-seq samples, respectively. The updated GeneFriends database contains co-expression data for 44,896 human genes and 31,236 mouse genes. The transcript co-expression data comprises of 145,455 human transcripts and 66,327 mouse transcripts. The biotype of genes and transcripts for both human and mouse data is given in Table 1. One of the unique features of GeneFriends co-expression database are its co-expression maps for non-coding genes like Long non-coding RNA (lncRNA) and microRNA (miRNA) which can be useful in providing the insights for regulatory mechanism of gene expression at both transcriptional and post-transcriptional level. The updated GeneFriends database have co-expression data for nearly 16,450 human and 6,436 mouse non-coding genes.

**Table 1.** The biotype of genes/transcripts present in GeneFriends co-expression database

| **Co-expression database** | **Number of Protein coding genes** | **Number of non-coding genes (%)** | **Others (%)** |
| --- | --- | --- | --- |
| Human Genes<br>(n=44896) | 19642 (43.8%) | 16450 (36.6%) | 8804 (19.6%) |
| Human Transcripts<br>(n=145455) | 89433 (61.5%) | 39776 (27.3%) | 16246 (11.2%) |
| Mouse Genes<br>(n=31236) | 19715 (63.1%) | 6436 (20.6%) | 5085 (16.3%) |
| Mouse Transcripts<br>(n=66327) | 42852 (64.6%) | 12928 (19.5%) | 10547 (15.9%) |

\*n=total number of genes present in the co-expression database, Others=Pseudogenes, TR (T-cell receptor genes), IG gene (Immunoglobulin genes)

We have also compared the top 5% of ten randomly selected human genes and their co-expression partners, which are present in both previous version (van Dam et al. 2015) and new updated version of GeneFriends (Table 2). The percentage of the average overlap between the ten genes was 30.5% with standard deviation of 4.97%. This difference between the two versions could be due to the difference in number of samples. The previous version was constructed from only 4133 RNA-seq samples as compared to the updated version which is based on 46475 samples. However, when we compared the functional enrichment of the top 5% co-expressed partners for some of these genes, the overlap was stronger suggesting that although the overlap between the co-expressed partners was low but overall they were associated with similar functional categories (Supplementary Data).

**Table 2.** Overlapping of co-expression partners between old and new GeneFriends co-expression database

| **Gene name** | **Overlap percentage (new data vs old data)** |
| --- | --- |
| BCL6 | 33 |
| BRCA1 | 35 |
| CDKN2A | 29 |
| FOXO3 | 24 |
| IGF1R | 28 |
| PARP1 | 40 |
| SIRT1 | 30 |
| TP53 | 27 |
| TUG1 | 25 |
| CDC6 | 34 |

Old data = GeneFriends older version by van Dam et al. (2015)
New data = Updated version of GeneFriends

## GENEFRIENDS GENE AND TRANSCRIPT DATA COMPARISON

To explore the differences between the gene and transcript co-expression maps in human and mouse co-expression database, we compared the median of Pearson correlation values for each gene/transcript with respect to its co-expression partners across the GeneFriends database. For transcripts, median of different transcripts of the same gene was calculated for doing comparison. A total of 34920 human and 25459 mouse genes and its transcripts were analysed. 78% of human and 70% of mouse genes had more than one transcript. While comparing the co-expression maps of human genes and transcripts, the overall co-expression values of genes were significantly higher than the co-expression values of transcripts (Figure 2A). The range of Pearson’s correlation coefficient values was widely distributed in genes encompassing both positive and negative values (Figure 2A and 2B). However, in case of transcripts they had lesser positive correlation values. This observation could be due to the fact that the transcript values are the median of different transcripts of the gene and different transcripts of same gene may have different trends of correlation. Similar trends were observed for the mouse co-expression database, where mouse genes had higher correlation coefficients than transcripts (Figure 2B), although the range of correlation coefficients were not as widely distributed as in humans (Figure 2B). These results indicated that different transcripts arising from the same gene are often expressed under different conditions and are most likely to play different roles in different processes or sometimes these transcripts may even be non-functional (Li et al. 2014).

**Figure 2.**
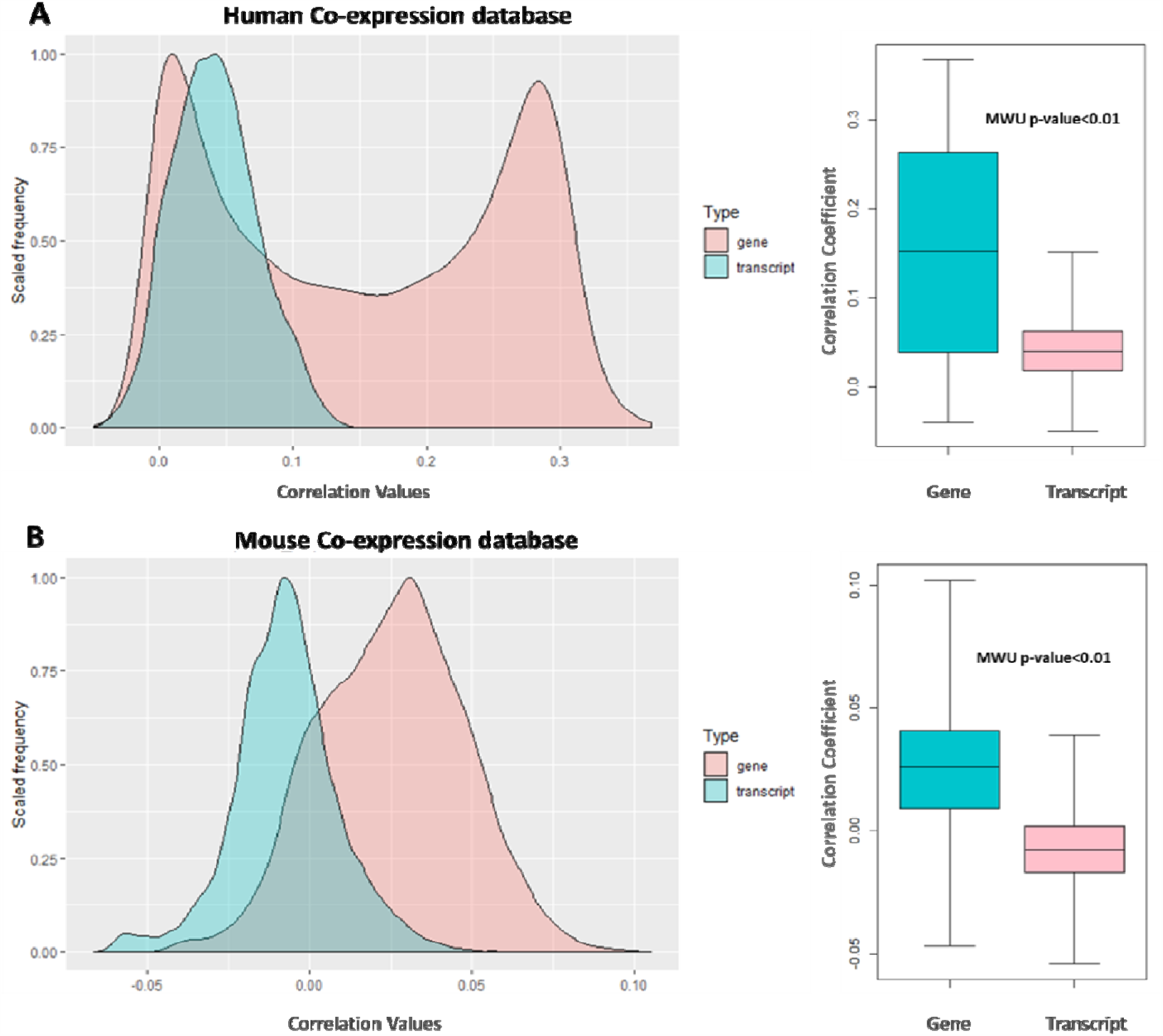
Comparison between human and mouse gene and transcript co-expression databases. (A) Comparison of correlation coefficient values between human genes and transcripts. (B) Comparison of correlation coefficient values between mouse genes and transcripts.

## PATHWAY ANALYSIS IN GENEFRIENDS

Since the primary purpose of the co-expression database is to determine the function of the co-expressed genes, we investigated the KEGG pathway genes to assess the consistency of the co-expression data with pathway annotations. We compared the number of enriched KEGG pathway genes between top and bottom 5% of co-expressed genes in human GeneFriends co-expression database. A total of 186 Kegg pathway gene sets from Molecular Signatures Database (MSigDB) v7.0 were analysed. The top 5% of co-expressed genes had significantly higher number [Median, Interquartile range (IQR) = 107(50-280)] of KEGG pathway enrichments in comparison to bottom 5% [Median (IQR) = 6(6-203)] (Supplementary Figure S1). This was followed by further analysing the top 5% of co-expressed genes with most enriched KEGG pathway genes for each 186 KEGG pathway gene sets (Supplementary Figure S2) and comparing top 20 and bottom 20 KEGG pathway annotations among human GeneFriends co-expression database (Figure 3A). The KEGG pathway enrichments like Glycolysis, insulin signalling, folate synthesis and WNT signalling were among the top 20 enriched KEGG pathway annotations. The annotations in top 20 were more or less related to metabolic pathways, DNA repair and signalling. The pathway annotations present in bottom 20 were associated with immune system and infection (Figure 3A). After this, we selected the top 20 genes from the GeneFriends database with maximum number of KEGG pathway annotations, and checked which pathways are most enriched in these top 20 genes (Figure 3B). Here also we observed that the pathways related to metabolism and cell signalling were among the top enriched KEGG pathways annotations. All these observations from KEGG pathway analysis indicated that genes that are enriched in KEGG pathway often tend to co-express together, underscoring that genes that are co-expressed tend to work cooperatively in the same biological pathways.

**Figure 3.**
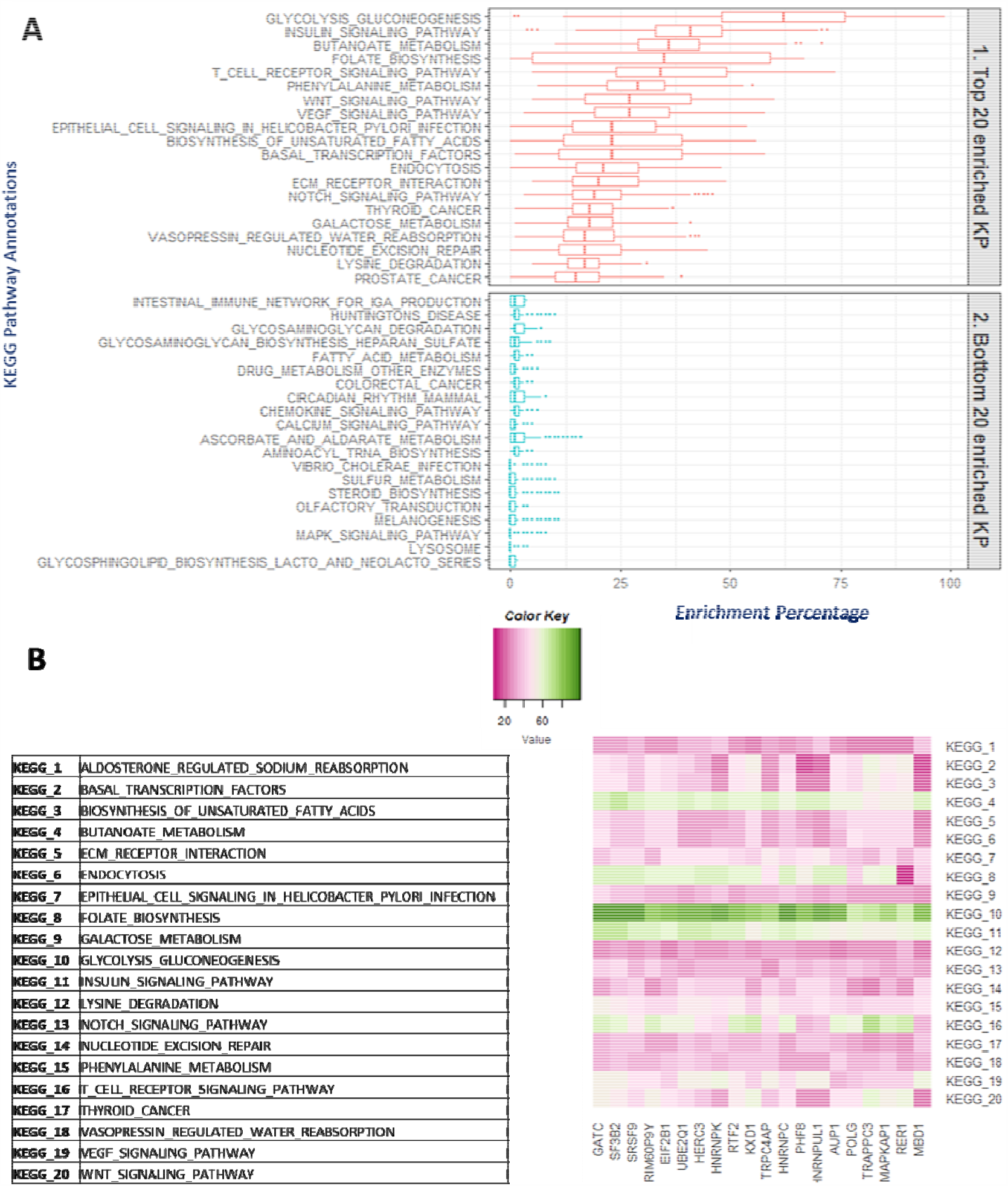
KEGG pathway enrichment analysis among the GeneFriends human database genes. (A) Top 20 and bottom 20 KEGG pathway annotations among the top 5% of GeneFriends human genes and their co-expressed partners. (B) KEGG pathway annotations among the top 20 GeneFriends genes with maximum number of KEGG pathway enrichments. The colour of the heat map represents the range of KEGG pathway enrichments among these 20 genes, Pink = low number of KEGG pathway enrichments and Green = High number of KEGG pathway enrichments.

**Figure 4.**
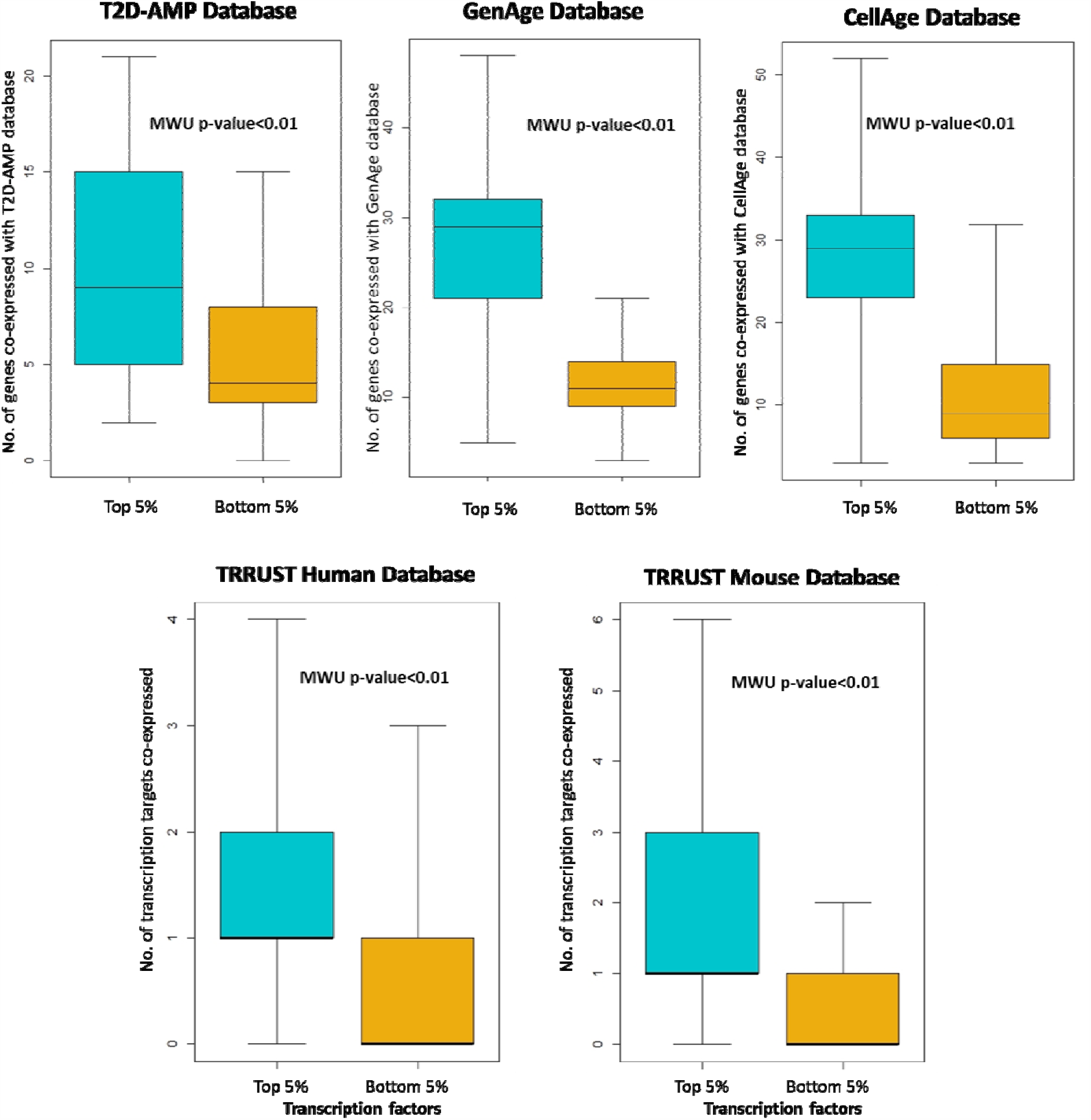
Comparing top and bottom 5% co-expressed gene partners of T2D-AMP, GenAge, CellAge and TRRUST database genes.

## VALIDATION OF GENEFRIENDS DATA

To assess the quality of the GeneFriends co-expression database we compared the top and bottom 5% of the genes that are present in some widely used databases. Genes from databases such as GenAge (Tacutu et al. 2018), CellAge (Avelar et al. 2020), T2D-AMP Knowledge Portal and TRRUST (Han et al. 2018) and their co-expressed partners were analysed to ascertain whether or not the genes that are linked to some diseases or processes tend to co-express together. GenAge is a curated database of genes related to ageing (Tacutu et al. 2018). We analysed co-expression data of 298 GenAge genes. The top 5% of GenAge genes present in GeneFriends database had significantly higher number of GenAge genes as their co-expressed partners as compared to the bottom 5% [Median(IQR): Top = 29(21-32); Bottom = 11(9-14)]. Similar trend was observed for 272 CellAge database (a curated database of cell senescence genes) and their co-expressed partners, where top 5% had significantly higher number of CellAge genes co-expressed in comparison to bottom 5% [Median(IQR): Top = 29(23-33); Bottom = 9(6-15)]. We were also interested to see how often genes that are related to some diseases may co-express with each other. To investigate this we analysed 132 type 2 diabetes (T2D) effecter genes from T2D-AMP database (https://t2d.hugeamp.org/gene/effectorGeneTable). We observed that T2D effector genes co-express with each other as top 5% had significantly higher number of T2D genes with respect to the bottom 5% [Median(IQR): Top = 9(5-15); Bottom = 4(3-8)].

To further validate our observations we also tested transcription factor and their targets from TRRUST database version 2 (Han et al. 2018). TRRUST database is a manually curated database of human and mouse transcriptional regulatory networks. As genes that co-express with each other may also help in co-regulating each other, hence we postulated that transcription targets should co-express with their respective transcription factors. We removed transcription factors where the relationship with the target was unknown. For human co-expression database, 603 human transcription factors were analysed. These transcription factors were then matched with 1710 transcriptional targets. The top 5% of co-expressed genes of all transcription factors had significantly higher number of transcriptional targets expressed in comparison to bottom 5% [Median(IQR): Top = 1(1-2); Bottom = 0(0-1)]. A total of 223 transcription factors had at least one transcriptional targets present in top 5% co-expressed genes. In case of mouse co-expression database, co-expression data for 703 mouse transcription factors were checked for 2100 transcriptional targets. Similarly for human transcription factors, top 5% mouse co-expression partners of transcription factors had significantly higher number of transcriptional targets in comparison to bottom 5% [Median(IQR): Top = 1(1-3); Bottom = 0(0-1)]. A total of 317 transcription factors had at least one transcriptional target present in top 5%. All these observations indicated that GeneFriends co-expression database is successfully able to identify the genes that are co-expressed and co-regulated together.

## COMPARISON OF HUMAN AND MOUSE CO-EXPRESSION NETWORKS

We analysed human and mouse co-expression networks from an updated GeneFriends co-expression database to decipher the evolutionary differences and similarities between human and mouse co-expression maps. We compared 24,434 genes that have a homolog in both human and mouse gene co-expression database. In our co-expression database, 14,911 genes were one-to-one orthologs, while the remaining mouse and human homologs had a one-to-many or many-to-many relationship. To understand the impact of duplication events on the divergence of humans and mice, we compared the dN/dS ratios of homologous genes with different types of homology (Fig. 5A). The one-to-one orthologs had the lowest dN/dS ratio as compared to the many to many, which had the highest dN/dS ratio. Next, we compared 14911 one to one orthologs among the top 5% of co-expressed genes. The dN/dS ratio values were divided into four groups to check how the increase/decrease in these values may relate to overlapping between two co-expression networks (Figure 5B). We observed that the group with the lowest dN/dS values had the highest number of overlapped co-expressed genes. This supported the hypothesis that non-synonymous substitutions influence the conservation of co-expression connectivity (Monaco et al. 2015). Therefore, the more the number of non-synonymous substitutions, the more conserved is a co-expression network.

**Figure 5.**
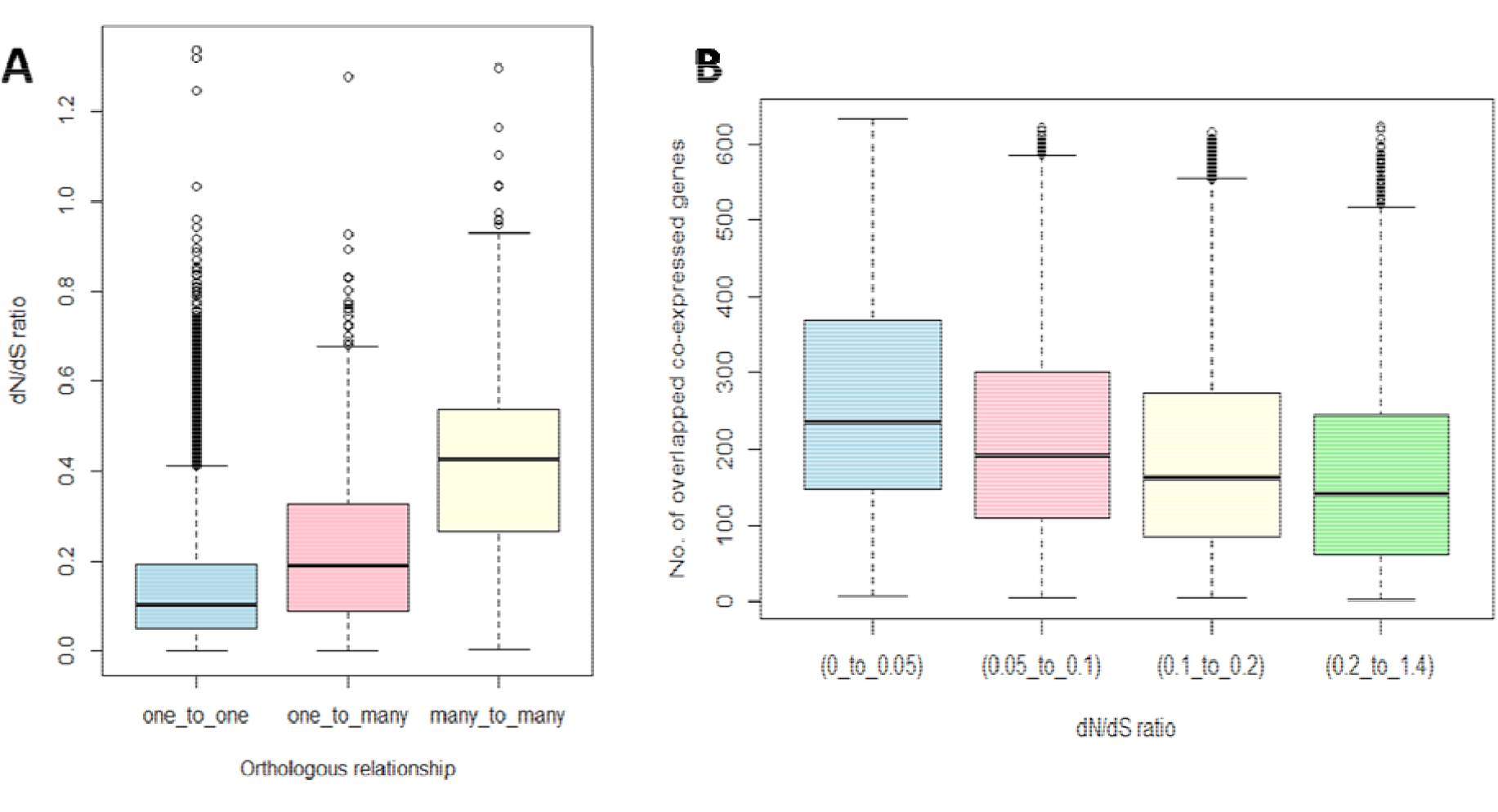
(A) Comparison of dN/dS values of homologs with three different relationships (One to one, one to many and many to many). The Mann-Whitney test showed significant difference between all three comparisons (One to one *vs* one to many, one to many *vs* many to many and one to one *vs* many to many). (B) Comparison of the dN/dS values between the top 5% of human and mouse co-expression gene networks.

## CONCLUSIONS

Large-scale gene co-expression networks have proven effective for analysing and discovering new gene functions and associations (Liesecke et al. 2019). With the large amounts of publicly available RNA-seq expression data, many co-expression databases are now using RNA-seq based data to generate co-expression networks. GeneFriends is unique in many ways, however, as compared to other available co-expression databases. The features that makes GeneFriends exceptional are its transcript based co-expression maps and inclusion of co-expression networks for non-coding genes. The GeneFriends database encompasses co-expression networks for about 16,000 and 6,000 non-coding genes for humans and mice, respectively. These transcripts and non-coding gene data based co-expression networks are crucial to understand the regulation of gene expression pattern, alternative splicing and dynamic regulation of transcripts in genome. Furthermore, we validated GeneFriends, and especially the results using curated transcription factor-transcriptional target database showed that genes that are co-expressed with each other also tend to co-regulate each other. Overall, in our latest version of human and mouse co-expression networks we hope to make GeneFriends more unique and valuable to the scientific community.

## METHODS

### Generation of co-expression database

Human RNA-seq read counts for 46475 samples were downloaded from the *recount2* database (Collado-Torres et al. 2017). Human gene expression data was downloaded with recount Bioconductor package (Collado-Torres et al. 2017) and transcript data was downloaded from recountNNLS R package (Fu et al. 2018). Mouse RNA-seq based read counts were obtained for 34322 samples from ARCHS4 database with rhdf5 Bioconductor R package (Lachmann et al. 2018). The human samples were aligned against the GRCh38 human reference genome, and mouse samples against the GRCm38 mouse reference genome. The reads were then normalized by dividing the expression per gene/transcript to the combined expression of all genes/transcripts per sample. Cancer-based studies were excluded to avoid any bias in the co-expression database moreover; cancer-related samples do not generalize well with overall co-expression networks.

To create co-expression maps we used weighted Pearson correlation method (van Dam et al. 2015). This was followed by constructing mutual rank maps by employing the same approach used in COXPRESdb (Obayashi et al. 2019). We used guilt by association method to create co-expression networks. The new GeneFriends database contains co-expression data for 44,896 human genes and 31,236 mouse genes. The transcript co-expression data comprises of 145,455 human transcripts and 66,327 mouse transcripts. The genes that were not expressed in at least 20% of the samples were excluded from the database. The biotype of genes and transcripts for both human and mouse data was identified using biomaRt.

### Functional and Pathway Analysis

We used WebGestalt (Liao et al. 2019) to do the Overrepresentation Enrichment Analysis for each of the gene ontology categories (Biological Process. Cellular Component and Molecular Function). The significance level was determined at FDR<0.05 and the multiple test adjustment was done using the Benjamini–Hochberg method. We verified our enrichment results by repeating the analysis using DAVID’s annotation clustering (Huang et al. 2009). p-value and FDR < 0.05 were considered significant. We also used *ClusterProfiler* Version 3.14.3 (Yu et al. 2012) to visualize the GO terms (FDR<0.05) obtained from DAVID. For KEGG annotation analysis, genes lists with their enriched KEGG pathway annotations were obtained from the Molecular Signature Database Version 6.2 (Subramanian et al. 2005; Liberzon et al. 2015). The box plot and heat map for KEGG pathway analysis were created using R.

### Evolution based Analysis

To identify any differences in the evolutionary conservation of genes present in human and mouse co-expression networks we performed dN/dS analysis. The dN/dS values were obtained from biomaRt release 96.

### Statistical Analysis

Mann-Whitney U tests was used to test the significance between the correlation coefficients among top 5% and bottom 5% co-expressed partners of genes and to compare the distribution of dN/dS scores between the human and mouse co-expression database. The median and Inter quartile ranges (IQR) were calculated by R package.

## ACKNOWLEDGEMENTS

This work and PR are supported by a Wellcome Trust (UK) research grant (Ref no:208375/Z/17/Z) to JPM. IL and ZF are supported by a BBSRC grant (BB/R014949/1) to JPM. KC is supported by a Mahidol-Liverpool Ph.D. scholarship from Mahidol University, Thailand, and the University of Liverpool, UK. We are also grateful for current and past members of the Integrative Genomics of Ageing Group for useful discussions, and in particular Sipko van Dam and Gianni Monaco.

## DATA AVAILABILITY

Human gene and transcript co-expression maps are available for download at http://www.genefriends.org

## SUPPLEMENTARY FIGURES

**Supplementary Figure S1.**
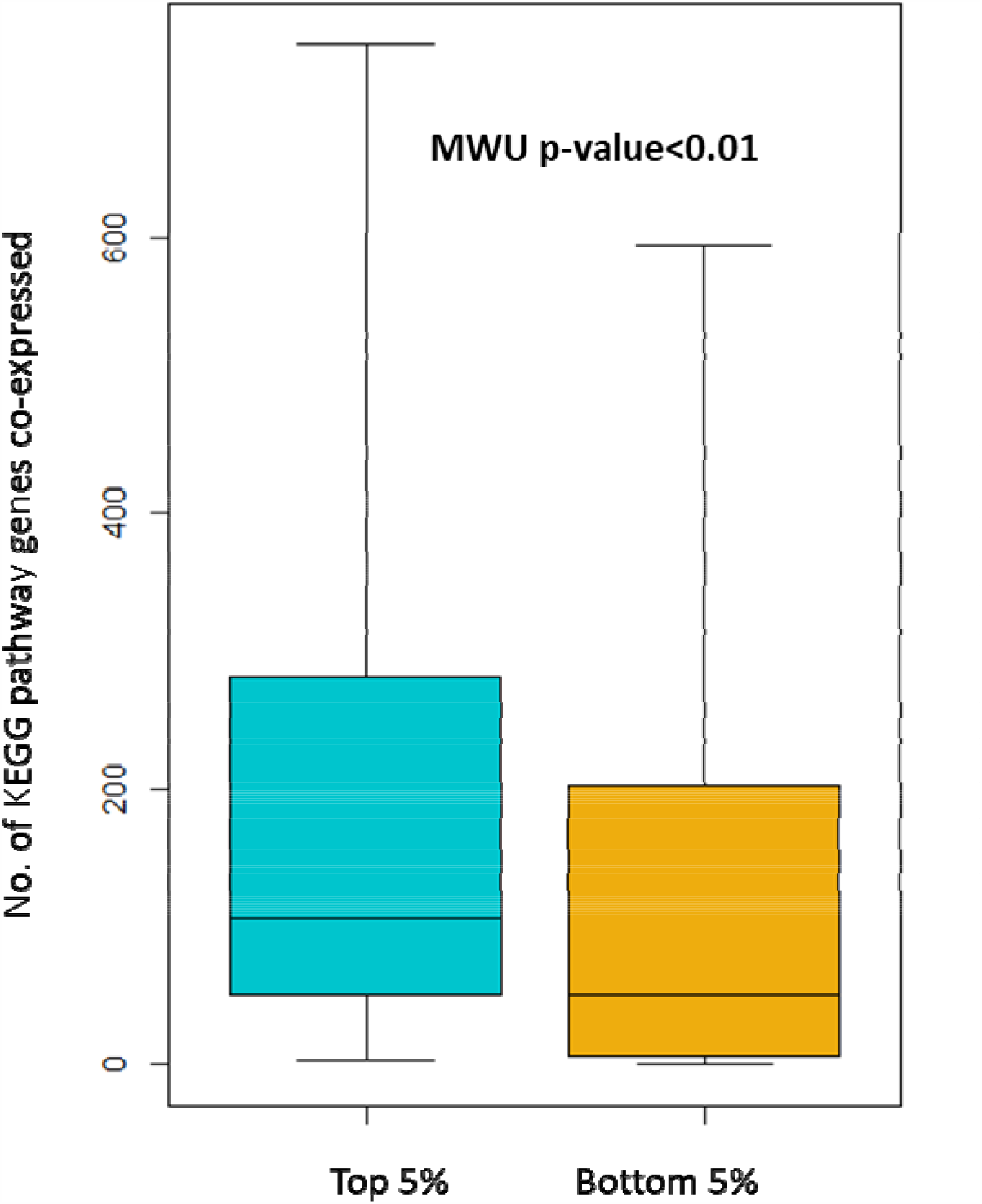
Comparison of KEGG pathway annotations among the top 5% and bottom 5% of the GeneFriends database genes.

**Supplementary Figure S2.**
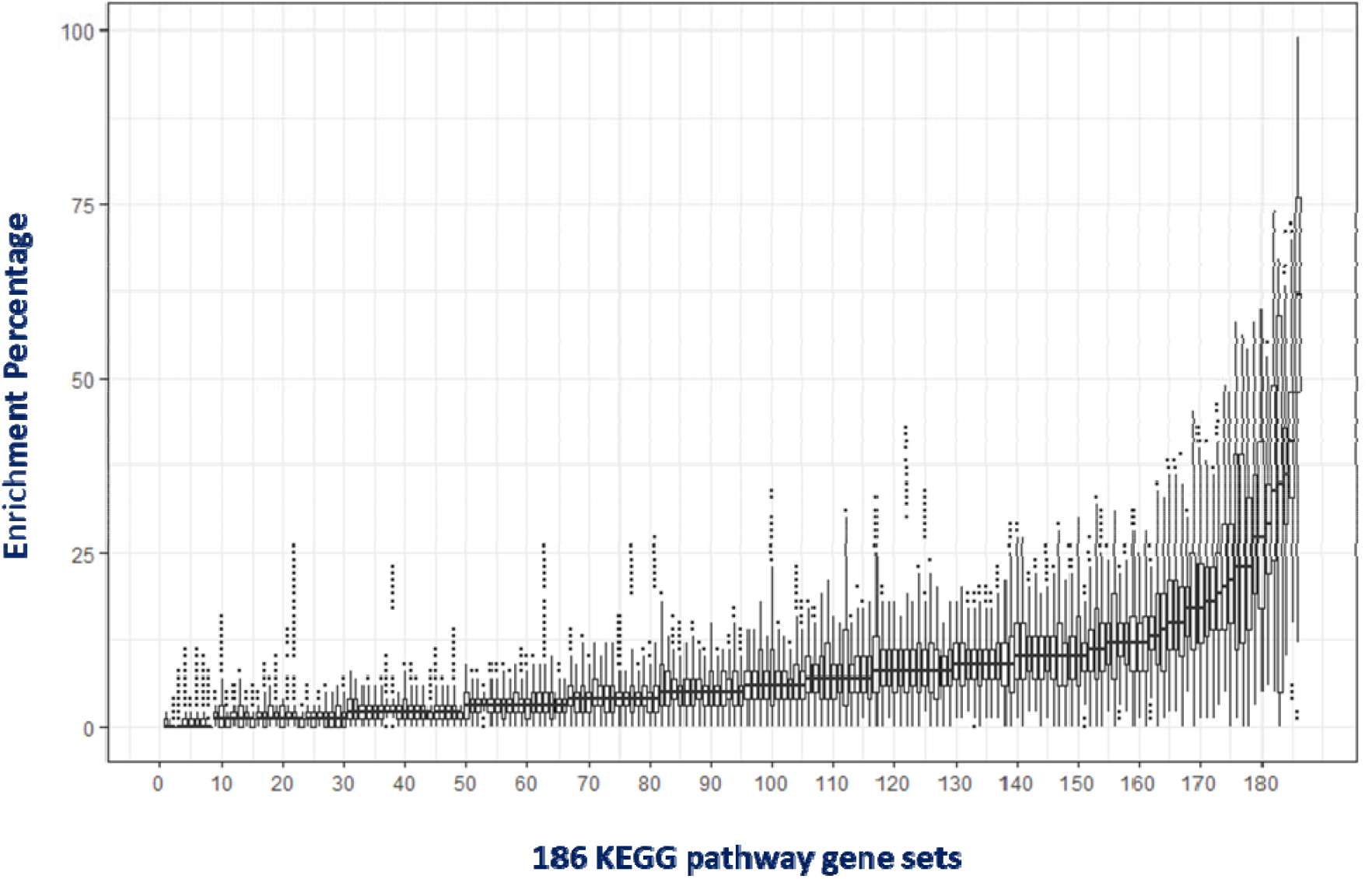
Comparison of 186 KEGG pathway annotations among top 5% of GeneFriends database genes.

